# Digital 3D Brain MRI Arterial Territories Atlas

**DOI:** 10.1101/2021.05.03.442478

**Authors:** Chin-Fu Liu, Johnny Hsu, Xin Xu, Ganghyun Kim, Shannon M. Sheppard, Erin L. Meier, Michael I. Miller, Argye E. Hillis, Andreia V. Faria

## Abstract

The locus and extent of brain damage in the event of vascular insult can be quantitatively established quickly and easily with vascular atlases. Although highly anticipated by clinicians and clinical researchers, no digital MRI arterial atlas is readily available for automated data analyses. We created a digital arterial territory atlas based on lesion distributions in 1,298 patients with acute stroke. The lesions were manually traced in the diffusion-weighted MRIs, binary stroke masks were mapped to a common space, probability maps of lesions were generated and the boundaries for each arterial territory was defined based on the ratio between probabilistic maps. The atlas contains the definition of four major supra- and infra-tentorial arterial territories: Anterior, Middle, Posterior Cerebral Arteries and Vertebro-Basilar, and sub-territories (thalamoperforating, lenticulostriate, basilar and cerebellar arterial territories), in two hierarchical levels. This study provides the first publicly-available, digital, 3D deformable atlas of arterial brain territories, which may serve as a valuable resource for large-scale, reproducible processing and analysis of brain MRIs of patients with stroke and other conditions.

## 1 BACKGROUND AND SUMMARY

Vascular atlases provide means to quickly establish the locus and extension of brain damage in the event of vascular insult^1^, can assist in stroke treatment planning and determining recovery prognosis^2, 3^, and can be used to inform future clinical trials of relevant scores (e.g., ASPECTS^4^). Research-wise, atlas-based analysis increases the statistical power and greatly improves the feasibility and reproducibility of lesion-based studies that usually require large scale data. Prior to the advent and use of modern neuroimaging techniques, vascular atlases were derived mainly from post-mortem injection studies^5–7^ and depicted schematic representations of the vascular territories. Such studies typically included single cases or very small samples, and thus, these atlases did not account for the high inter-individual variability in vascular distributions. Two-dimensional, manually-drawn atlases^8–12^ also suffer from a lack of flexibility; if the template slices provided in the atlas do not match the slice thickness or orientation of acquired clinical neuroimages, the atlas utility is diminished^1, 6, 9, 13, 14^. Even with well-aligned slices, printed templates lack precision and do not allow for quantitative assessment of vascular territories in a single patient.

Contemporary, digital atlases often derived from in vivo perfusion studies in healthy individuals^14–16^ or analysis of stroke lesions^1, 13, 17–19^ overcome the aforementioned limitations and can provide a fast and objective measure of vascular damage in individual patients. In particular, digital atlases that incorporate data from a large number of stroke patients are likely most representative of the vascular anatomy of older adults who are most likely to experience infarct. Recently, Kim et al.^18^ used a large dataset of stroke patients from academic centers in Korea to map supratentorial cerebral arterial territories. Despite its strengths, infratentorial and deep perforating arterial territories were not defined, even though approximately 20% of all strokes occur in both the infratentorial^20^ and deep perforating arterial zones^21^. Furthermore, a publicly accessible electronic version of the atlas is not available, thus reducing applicability. Wang et al.^22^ later used MRIs of 458 patients to create a “stroke atlas” using voxel-wise data-driven hierarchical density clustering. Although this atlas represents major and end-arterial supply territories, the difficulties in accessing the stability of small clusters and the lack of unitary-defined boundaries reduces applicability for individual brain mapping.

To remedy the limitations of existing vascular atlases, we created a novel whole-brain arterial territory atlas based on lesion distributions in 1,298 patients with acute stroke. Our atlas covers supra- and infra-tentorial regions, includes straightforward and robustly-defined watershed border zones, and contains hierarchical segmentation levels created by a fusion of vascular and classical anatomical criteria. The atlas was developed using diffusion weighted MRIs (DWIs) of patients with acute and early subacute ischemic strokes, expert annotation (lesion delineation), and metadata (radiological evaluation) of a large and heterogeneous sample, both in biological terms (e.g.., lesion pattern, population phenotypes) and technical terms (e.g., differences in MRI protocols and scanners). The biological and technical heterogeneity of the sample is advantageous in that it increases the generalization and robustness of the atlas, and consequently its utility. This first publicly-available, digital, 3D deformable atlas can be readily used by the clinical and research communities, enabling to explore large-scale data automatically and with high levels of reproducibility.

## 2 METHODS

### 2.1 Cohort

This study included Magnetic Resonance Images (MRIs) of patients admitted to the Johns Hopkins Stroke Center with the clinical diagnosis of acute stroke, between 2009 and 2019. We have complied with all relevant ethical regulations and the Johns Hopkins Institutional Board Review guidelines for using this image archive. Baseline MRIs adequate for clinical analysis and evidence of ischemic stroke in DWI were included (please see the flowchart of data inclusion in Figure 1). From the 1,878 patients identified under these criteria, 580 individuals were excluded because their stroke was 1) of confirmed cardioembolic origin, 2) bilateral, 3) exclusively within “watershed” areas, or 4) multifocal / involving more than one major arterial territory, per radiological evaluation of the MRI. Ultimately, 1,298 MRIs were included with acute or early subacute ischemic strokes affecting exclusively one major arterial territory (anterior, middle, and posterior cerebral, or vertebro-basilar arteries).

**Figure 1.**
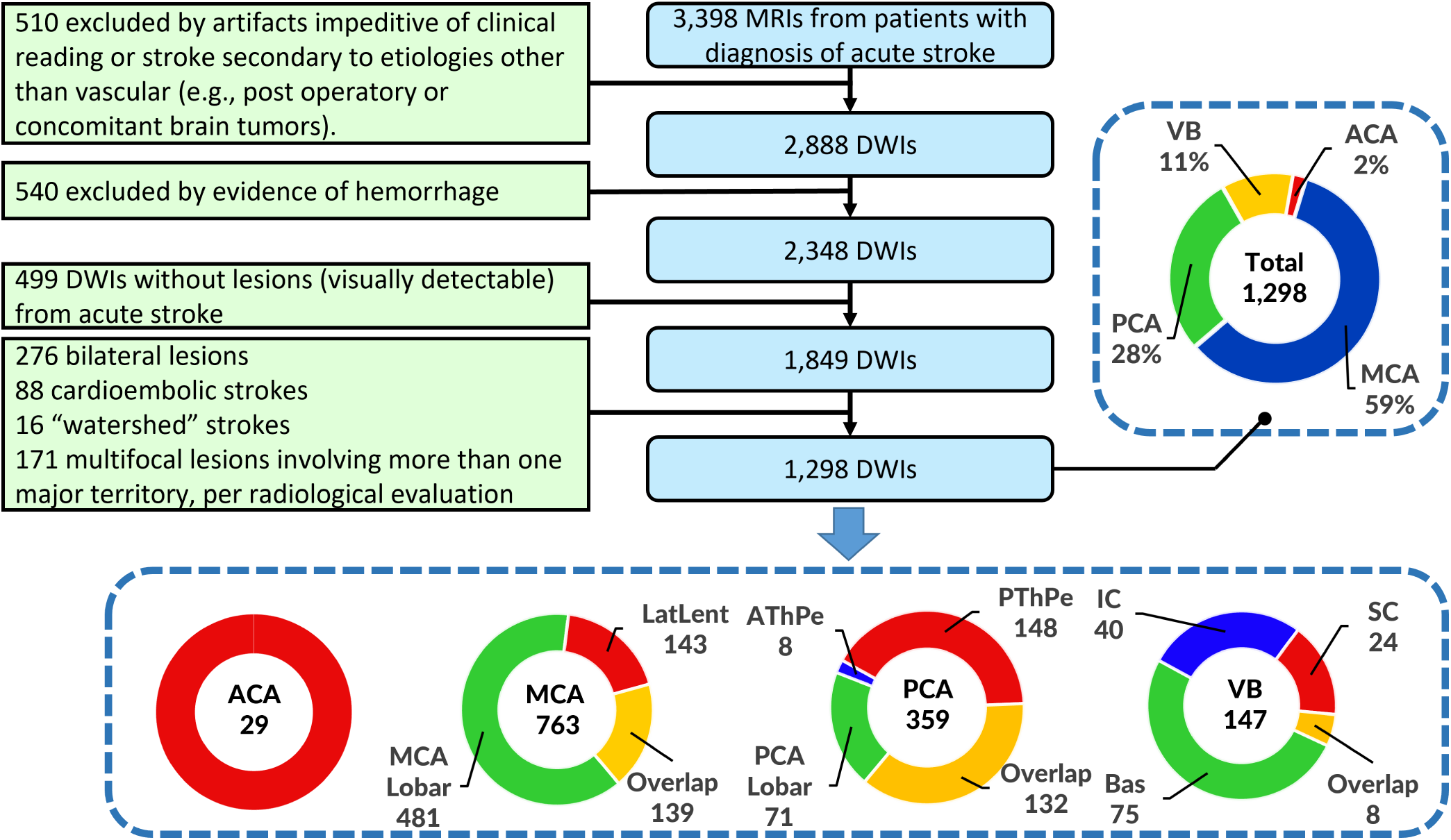
Flowchart of data inclusion. Abbreviations: anterior, middle, and posterior cerebral arteries (ACA, MCA, PCA, respectively), vertebro-basilar (VB), lateral lenticulostriate (LatLent), posterior and anterior thalamo-perforators (PThPe, AThPe), inferior and superior cerebellar (IC, SC), basilar (Bas)

The characteristics of the cohort are summarized in Table 1. The distribution of strokes according to arterial territories (MCA > PCA > VB > ACA) and the demographic and clinical characteristics reflect the general population of stroke patients. We note that our population includes a higher percentage of Black, and lower percentage of Hispanic/Latinx and Asian patients than many urban stroke centers, therefore a regional bias may exist. The vast majority of scans were performed more than 2 hours after the onset of symptoms, reducing the odds of significant further change in the lesion volume^23^.

**Table 1.** Population, lesion and scanner profiles per Vascular Territory. ACA, PCA, MCA stand for Anterior, Posterior, and Middle Cerebral Artery territories, VB stands for Vertebro-Basilar territory. Statistical significant differences at p-value <0.05 are marked with “*”; Continued variables showed as mean ± standard deviation [maximum, minimum]; missing data

| **Vascular Territories** | **Total** | **ACA** | **MCA** | **PCA** | **VB** |
| --- | --- | --- | --- | --- | --- |
| **Number of DWIs** | 1298 | 29 | 763 | 359 | 147 |
| **Age in years** | 62.3 ± 14.6<br>[99, 18] | 65.8 ± 15.7<br>[85, 22] | 63.0 ± 15.1<br>[99, 18] | 61.4 ± 13.5<br>[96, 20] | 60.4 ± 14.4<br>[93, 21] |
| **Sex** |  |  |  |  |  |
| Male | 686 (52.9%) | 13 (44.8%) | 394 (51.6%) | 197 (54.9%) | 82 (55.8%) |
| Female | 612 (47.1%) | 16 (55.2%) | 369 (48.4%) | 162 (45.1%) | 65 (44.2%) |
| **Race** |  |  |  |  |  |
| African American | 600 (62.6%) | 16 (69.6%) | 316 (58.7%) | 190 (69.3%) | 78 (63.4%) |
| Caucasian | 334 (34.9%) | 7 (30.4%) | 208 (38.7%) | 78 (28.5%) | 41 (33.3%) |
| Asian | 24 (2.5%) | 0 | 14 (2.6%) | 6 (2.2%) | 4 (3.3%) |
| Missing data | 340 | 6 | 225 | 85 | 24 |
| **NIHSS** | 5.4 ± 5.9<br>[31, 0]; 551 | 3.4 ± 5.4<br>[19, 0]; 10 | * 7.1 ± 6.8<br>[31, 0]; 325 | 3.0 ± 2.7<br>[14, 0]; 150 | 2.9 ± 3.8<br>[22, 0]; 66 |
| **Days of hospital stay** | 5.2 ± 5.5<br>[39, 0]; 416 | 4.8 ± 3.3<br>[13, 1]; 8 | 5.9 ± 5.7<br>[39, 0]; 260 | * 3.6 ± 4.3<br>[38, 0]; 107 | 5.5 ± 6.5<br>[34, 0]; 41 |
| **Onset to MRI in hours** |  |  |  |  |  |
| <2 | 79 (9.5%) | 4 (19.0%) | 52 (11.1%) | 18 (7.7%) | 5 (4.8%) |
| 2-6 | 170 (20.5%) | 5 (23.8%) | 90 (19.1%) | 46 (19.7%) | 29 (27.9%) |
| 6-12 | 166 (20.0%) | 6 (28.6%) | 87 (18.5%) | 56 (23.9%) | 17 (16.3%) |
| 12-24 | 346 (41.7%) | 4 (19.0%) | 203 (43.2%) | 96 (41.0%) | 43 (41.3%) |
| >24 | 68 (8.2%); | 2 (9.5%); | 38 (8.1%); | 18 (7.7%) | 10 (9.6%); |
| Missing data | 469 | 8 | 293 | 125 | 43 |
| **MRI magnetic field** |  |  |  |  |  |
| 1.5T | 869 (66.9%) | 16 (55.2%) | 517 (67.9%) | 238 (66%) | 98 (66.7%) |
| 3T | 429 (33.1%) | 13 (44.8%) | 246 (32.1%) | 121 (34%) | 9 (33.3%) |
| **Lesioned hemisphere** |  |  |  |  |  |
| Left | 682 (52.5%) | 23 (79.3%) | 385 (50.5%) | 189 (52.6%) | 85 (57.8%) |
| Right | 616 (47.5%) | 6 (20.7%) | 378 (49.5%) | 170 (47.4%) | 62 (42.2%) |
| **Stroke volume in ml** | 21.6 ± 47.1<br>[403.5, 0.028] | 7.6 ± 10.9<br>[44.4, 0.15] | * 32.2 ± 58.1<br>[403.5, 0.06] | 6.4 ± 14.0<br>[131.6, 0.028] | 6.8 ± 13.8<br>[70.9, 0.084] |
| **Relative stroke volume** |  |  |  |  |  |
| % of arterial territory | 7.8% | 5.5% | 9.2% | 5.5% | 6.9% |
| % of brain volume | 1.50% | 0.58% | 2.24% | 0.43% | 0.45% |

### 2.2 Image Analysis

MRIs were obtained on seven scanners from four different vendors, in different magnetic fields, with dozens of protocols. The DWIs had high in plane (axial) resolution (1×1mm, or less), and typical clinical large slice thickness (ranging from 3 to 6 mm). The delineation of the stroke core was defined in the DWI by 2 experienced evaluators and revised by a neuroradiologist until reaching a final decision by consensus. Details are described in Technical Validation. The binary lesion masks (ischemic lesion = 1; brain and background = 0) were then used to create probabilistic maps, as shown in Figure 2.

**Figure 2.**
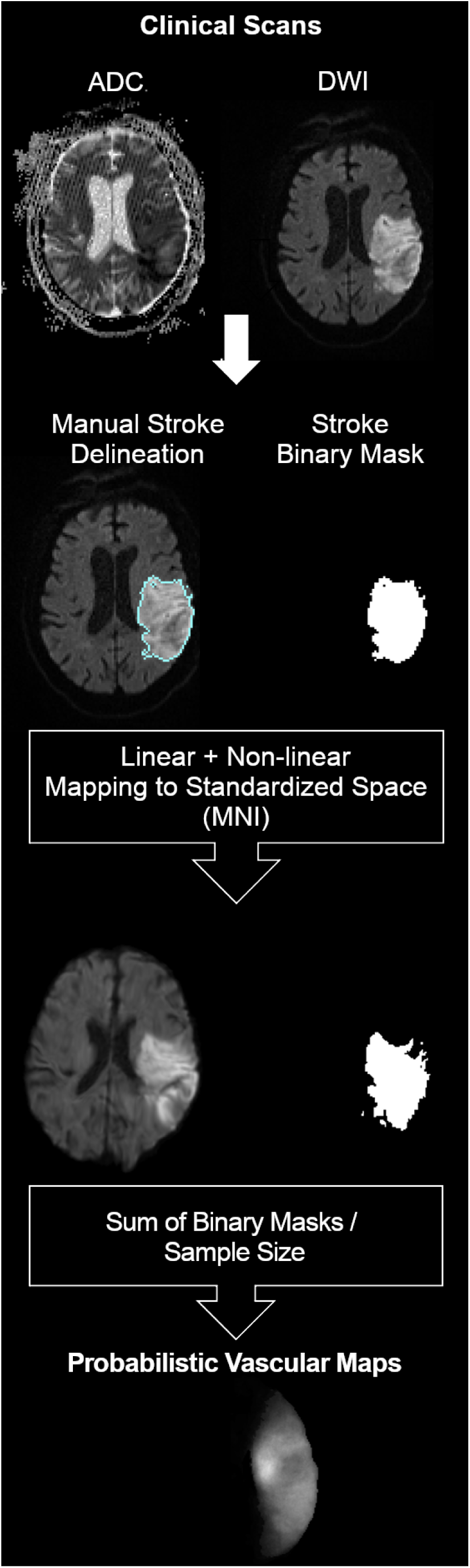
Schematic representation of the imaging processing.

Creating probabilistic maps and atlases requires transforming images to a common space for consistency among image coordinates and biological structures. Clinical stroke images offer three additional challenges to image mapping: 1) the typical high slice thickness, often associated with large rotations out-of-plane, 2) the presence of drastic changes in image morphology and intensity, caused by the stroke as well as associated chronic conditions (e.g., white matter microvascular chronic lesions), and 3) the variable and often remarkable degree of brain atrophy in elderly participants who primarily constitute the stroke population. As linear deformations are not enough to accurately map internal brain shapes, and high elastic transformations (e.g., diffeomorphic) are very sensitive to focal abnormalities in morphology and contrast, we applied sequential steps of linear and non-linear deformation, using Broccoli^24^. To minimize the effects of the acute stroke, we used the “B0” images (less affected by the acute lesion intensity) for mapping. As a template, we used the JHU_SS_MNI_B0^25^, a subject in standard MNI coordinates. The optimization of the parameters and other details of the image mapping are described in the Technical Validation.

### 2.3 Arterial territory maps: average method

The cohort was divided into four groups based on the major arterial territory that was exclusively affected in each patient, according to radiological classification: 1) Anterior Cerebral Artery (ACA), 2) Middle Cerebral Artery (MCA), 3) Posterior Cerebral Artery (PCA), or 4) Vertebro-Basilar artery (VB). The MCA infarcts affected the lateral lenticulostriate territory, or the “lobar” (cortex and adjacent subcortical white matter) MCA territory, or both; the PCA infarcts affected the posterior and anterior thalamoperforating territory, or the “lobar” PCA territory, or a combination of both; the ACA infarcts affected the medial lenticulostriate territory, or the “lobar” ACA territory, or both; the VB infarcts affected the inferior or superior cerebellar areas, or the basilar territory, or a combination of both.

Probabilistic maps were generated by summing the lesion binary mask in the common space and dividing by the sample size for each of these territories. Given an annotated pair (*s*_*i*_, *X*_*i*_) for each subject, where *s*_*i*_ is the group label, 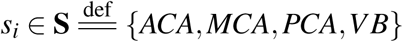, and *X*_*i*_ is the lesion mask, *i* is the index of subjects. Let *X*_*i, j*_ denote the voxel *j* in the lesion mask *X*_*i*_, where *j* is the index of voxels and *N*_*s*_ denote the sample size of the *s* group, ∀*s* ∈ **S**. *X*_*i, j*_ = 1, if *X*_*i, j*_ is a lesioned voxel; otherwise, it is 0. The likelihood of each voxel *X*_*i, j*_ belonging to *s* group can be estimated as

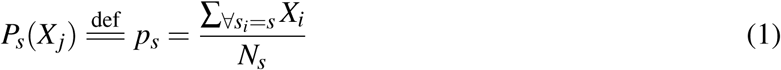

, for *s* ∈ **S**. For example,

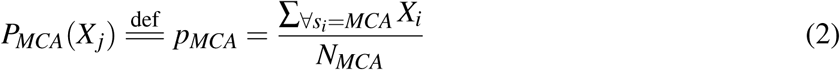

denotes the probabilistic map *p*_*MCA*_ for MCA territory. Similarly, we compute *p*_*ACA*_, *p*_*PCA*_, and *p*_*VB*_. These probabilistic maps (Figure 3) can be considered as an estimate for the likelihood of voxels belonging to a certain territory. The more samples in a group, the more accurate the estimate is.

**Figure 3.**
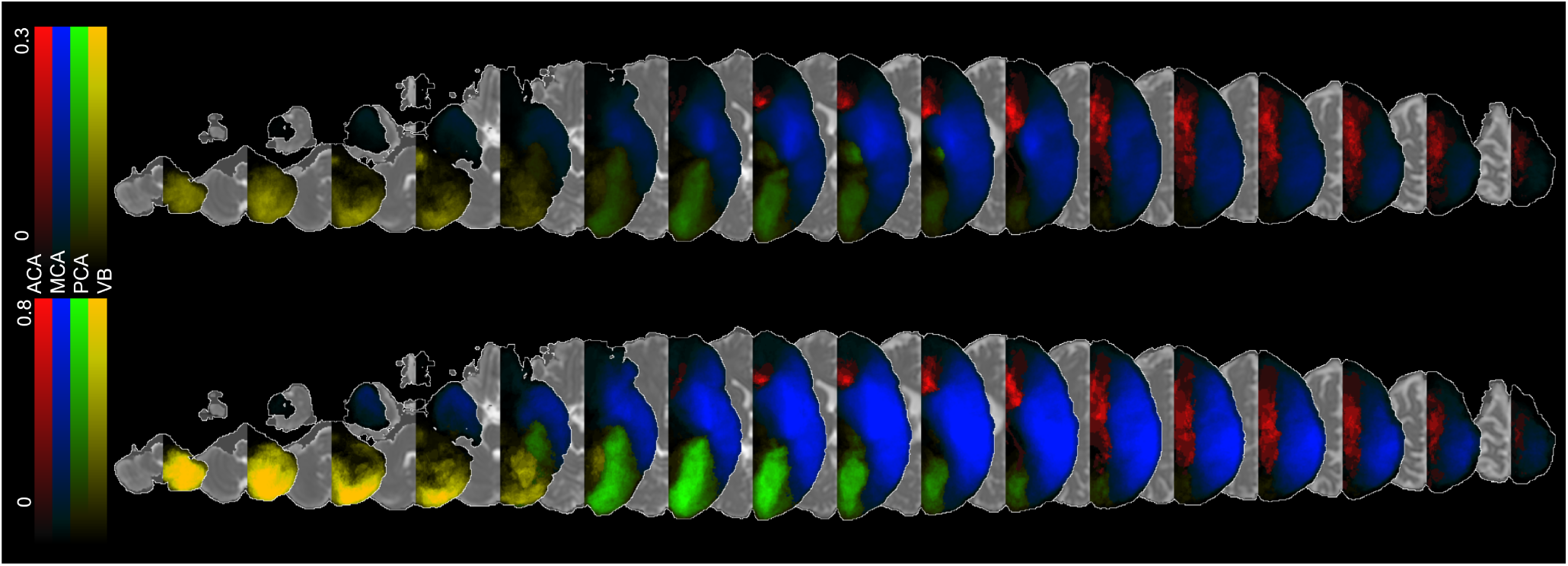
Probabilistic maps of the four major arterial territories: anterior cerebral artery (ACA), medium cerebral artery (MCA), posterior cerebral artery (PCA), and vertebro-basilar (VB), calculate via averaging the lesions masks (top) or via the BMM-method (bottom).

For the “border zone” between the major cerebral arteries (ACA, MCA, PCA), we considered each voxel that had a probability greater than zero to belong to two arterial territories. By dividing the probabilities, we estimated the likelihood that voxels in the border zone belong to one or another major arterial territory (Figure 7). For example, a ratio of *p*_*MCA*_/*p*_*PCA*_ > 1 indicates that the given “border zone” voxel is more often affected in MCA strokes than in PCA strokes while *p*_*MCA*_/*p*_*PCA*_ <1 indicates the opposite. We acknowledge the sample imbalance (the ACA group is smaller than the others) as a possible bias for the conditional probability ratio.

### 2.4 Arterial territory maps: Bernoulli Mixture Model (BMM) method

The BMM method^26^ implicitly captures potential voxel-level correlations between spatial and volumetric features of lesion maps by generating probabilistic maps of each territory via the cluster of lesions with the most similar volumes and spatial features. Assume that for each region of interest, ROI (i.e. ACA, MCA, PCA, and VB), their lesion maps *X*_*i*_ can be generated by a mixuture of *K* clusters. The voxel *X*_*j*_ of the *k* cluster is binary as {0, 1} for non-lesioned or lesioned voxel with the Bernoulli distribution of {*µ*_*k, j*_}, where *i* is the index of subjects/samples, *j* is the index of voxels, and *k* is the index of clusters. And each cluster’s prior probability is denoted as *π*_*k*_. Therefore, the likelihood of each ROI can be represented as

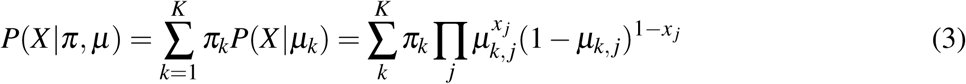

Given *π* and *µ*, the expectation and covariance matrix of *X, E*[*X*] and *Cov*[*X*], can be calculated as

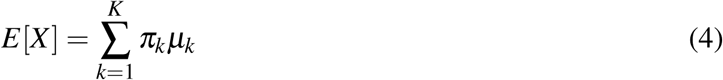

and

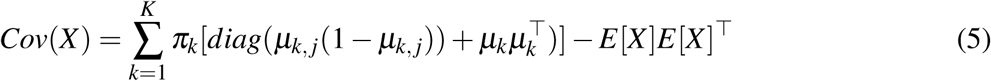

, where the *diag*(*µ*_*k, j*_(1 − *µ*_*k, j*_)) represents the diagonal matrix whose diagonal entries equal to *µ*_*k, j*_(1 − *µ*_*k, j*_). Since *Cov*(*X*) is not a diagonal matrix, the correlations between voxels could be captured by this model once *µ* and *π* are determined.

Now, assume *N* subjects of stroke maps *X*_*i*_, {*X*_1_, *X*_2_, …, *X*_*N*_} are generated independently, the log likelihood function of all subjects is

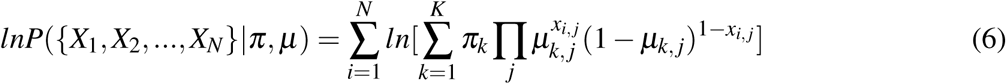

, where *x*_*i, j*_ denotes the voxel *j* in the lesion map of the subject *i*.

Our goal is to find the best *π* and *µ* to maximize the log likelihood function for given subjects of stroke maps *X*_*i*_, {*X*_1_, *X*_2_, …, *X*_*N*_}. As^26^, we applied the Expectation-Maximization (EM) algorithm to find the optimal *π* and *µ*, by introducing indicators *z*_*i,k*_ to get the complete-data log-likelihood function as

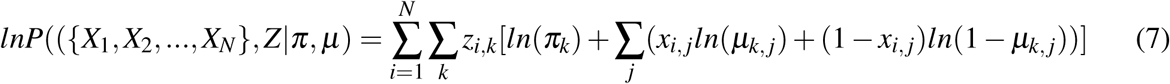

, where *Z* = {*z*_*i,k*_} for all *i* and *k*.

Therefore, the expectation of the complete-data log-likelihood function over *Z* is

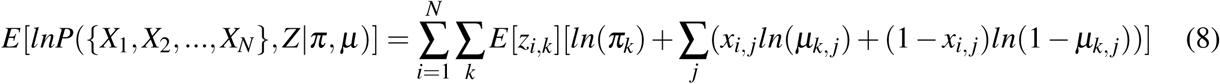

Then, by applying EM algorithm, we can get

1. E-step:

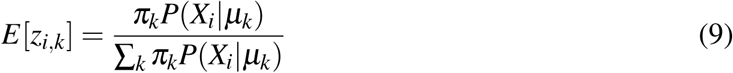
2. M-step:

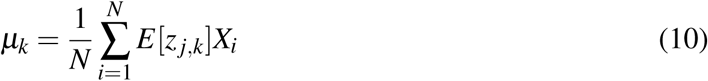

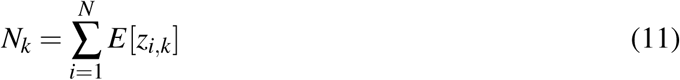

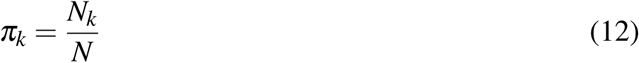

We can now calculate the *µ*_*k*_ map for the cluster *k* in a certain ROI. If the total number of clusters is 1, *K* = 1, the *µ* equals to the likelihood map *p*_*s*_ as calculated by the average method, Equation (1), which maximizes its corresponding log likelihood function for given *N*_*s*_ subjects of stroke maps *X*_*i*_, 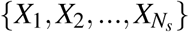 for each *s* ∈ 𝕊. If the number of clusters is more than one, the likelihood of a voxel *X*_*j*_ to belong to a certain ROI can be estimated by the cluster in which lesions have similar volume and spatial features. Consequently, this cluster is likely to be a better representation for such voxel. Therefore, we calculate the final *P*_*s*_(*X*_*j*_) as

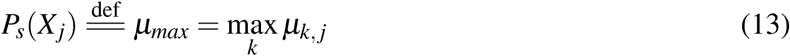

In addition, the expectation of the lesion volume for each cluster *k, E*[*V*_*k*_], can be estimated by

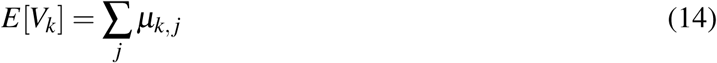

and the expectation of the number of subjects in cluster *k* is

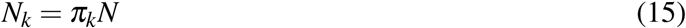

The remaining question is how to determine the number of clusters *K* for each ROI. If *K* is too large, we might define *P*_*s*_(*X*_*j*_) by a too small group of sample. Therefore, we chose the largest *K* for each BMM cluster contains at least 10% of the subjects in its ROI group. By doing so, *K* = 2 was chosen for *ACA, K* = 4 was chosen for *MCA, K* = 3 was chosen for *PCA*, and *K* = 3 was chosen for *VB*. The representation of each of these clusters is shown in Technical Validation.

The clusters *µ*_*max*_ were calculated to generate the new vascular territory probabilistic map *P*_*s*_(*X*_*j*_). Compared to average method, the BMM probability maps show higher and more uniformly distributed probabilities (Figure 3). Take MCA as an example: while the average method has higher values in the neighborhood of the basal ganglia (the maximum probability is about 0.3), the BMM probabilistic maps have higher and more similar values widespread over the MCA territory. This is due to the fact that with BMM, the likelihood of a voxel *X*_*j*_ to belong to a certain ROI can be estimated by the cluster in which lesions have similar volume and spatial features.

As for the border zone, we first determined the voxels in BMM probability maps, *µ*_*max*_, with high probability to be classified as “lesioned”. Due to the assumption that the stroke affects one exclusive major arterial territory in each case, a voxel with high probability to be a “lesioned voxel” has also a high probability to be within that major arterial territory. Conversely, a voxel classified as “non-lesioned” is unlikely to be within that territory. Hence, given *µ*_*k*_ from BMM for a certain ROI, we calculated the variance of each voxel *j* for each cluster by *µ*_*k, j*_(1 − *µ*_*k, j*_), because each voxel in each cluster has Bernoulli distribution of *µ*_*k, j*_. A voxel j is classified as 0 (non-lesioned voxel) within a certain ROI if

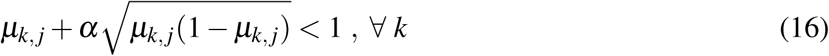

, where *α* is the parameter to determine the confidence level. On the other hand, a voxel j is classified as 1 (lesioned voxel) within a certain ROI if at least one *µ*_*k*_ makes

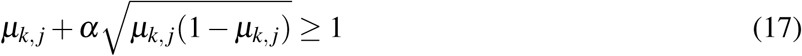

If a voxel is claimed by more than one territory, the one with the largest *µ*_*max*_ “wins” that voxel. The result of this process is illustrated in Figure 4. Regions less stroke-prone and territories with small samples (e.g., ACA) had, understandably, more voxels that were not “claimed” by any territory. These voxels were eventually labeled by water-spreading to the closest territory.

**Figure 4.**
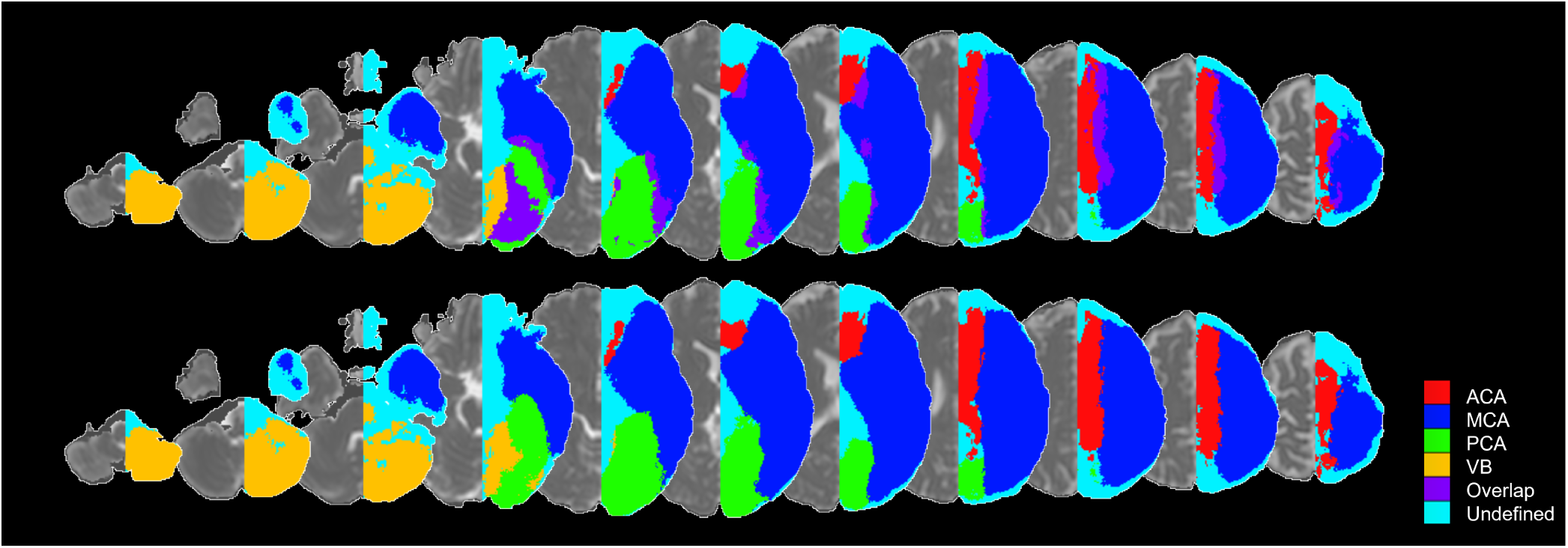
“Territory-claimed” voxel maps (*α* = 3). The colors represent the major arterial territories, as well as voxels claimed by more than one territory (“overlap”) and by none (“undefined”). The bottom row is after the assignment of the “overlap” voxels to a major territory.

### 2.5 Definition of Arterial Territories

Our ultimate goal was to create a 3D deformable atlas of the major arterial territories, which requires sharp boundaries between neighbor regions. After adding the major probabilistic maps (ACA, MCA, PCA, VB) obtained with the average method, the vast majority of voxels belonged to an exclusive vascular territory and were attributed an integer intensity that identified that territory. When a voxel belonged to more than one vascular territory (which occurred in the border zone) it was attributed to the territory with the highest probability. Discontinuities in the border zone were approximated by a straight line.

The sharp boundaires between neighbor regions were validated using the BMM-derived “territory-claimed” voxel maps, as that shown in 4. The “border zone” as defined by the probability ratios or by the BMM “territory-claimed” voxel maps highly agree, as shown in Figure 5, and very minor adjusts were made to conciliate them. The resulting atlas had a relatively coarse contour and was smoothed by a kernel filter followed by manual editing by an expert neuroradiologist. The manual adjusts were “cosmetic” (no more than 3 voxels, in plane, at 1×1×1 mm resolution).

**Figure 5.**
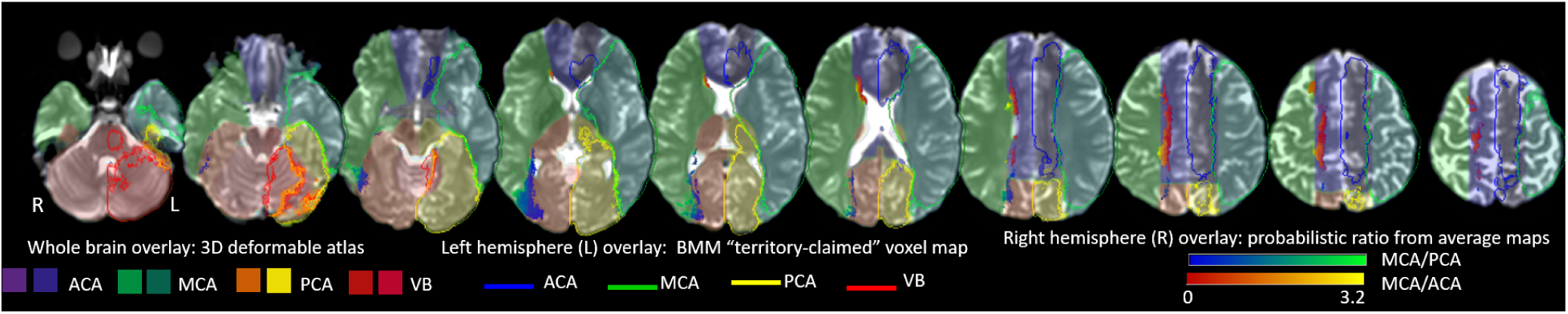
Representation of the border zone as defined by the different methods. Over the right hemisphere (R): probability ratios from the average maps (for better visualization of the probability ratios, see Figure 7D). Over the left hemisphere (L): limits of a BMM “territory-claimed” voxel map. Over the whole brain, as translucent colors: our digital (sharp-boundaries) atlas of major vascular territories.

The same procedure, based on probability ratios, was applied to define the sub-territories of lateral lenticulostriate (within MCA), thalamoperforating (within PCA), and anterior and posterior cerebellar and basilar (within VB) arteries. The “lobar” sub-areas of MCA (frontal, parietal, temporal, insular, occipital) and PCA (occipital, temporal), were defined based on classic anatomical landmarks (gyri, as defined in our previous atlas^27^). In less stroke-prone territories, such as medial lenticulostriate and anterior thalamopeforating (that have less than 10 cases of exclusive lesions in each), the probabilistic maps would be less reliable and we chose to rely on prior anatomical knowledge. The medial lenticulostriate territory, within ACA, corresponds to the topography of corpus callosum. The anterior thalamoperforating territory, within PCA, corresponds to the topography of the mesial temporal area and hippocampus. As a result, we created a hierarchical (2-level) brain atlas of arterial territories (Figure 6).

**Figure 6.**
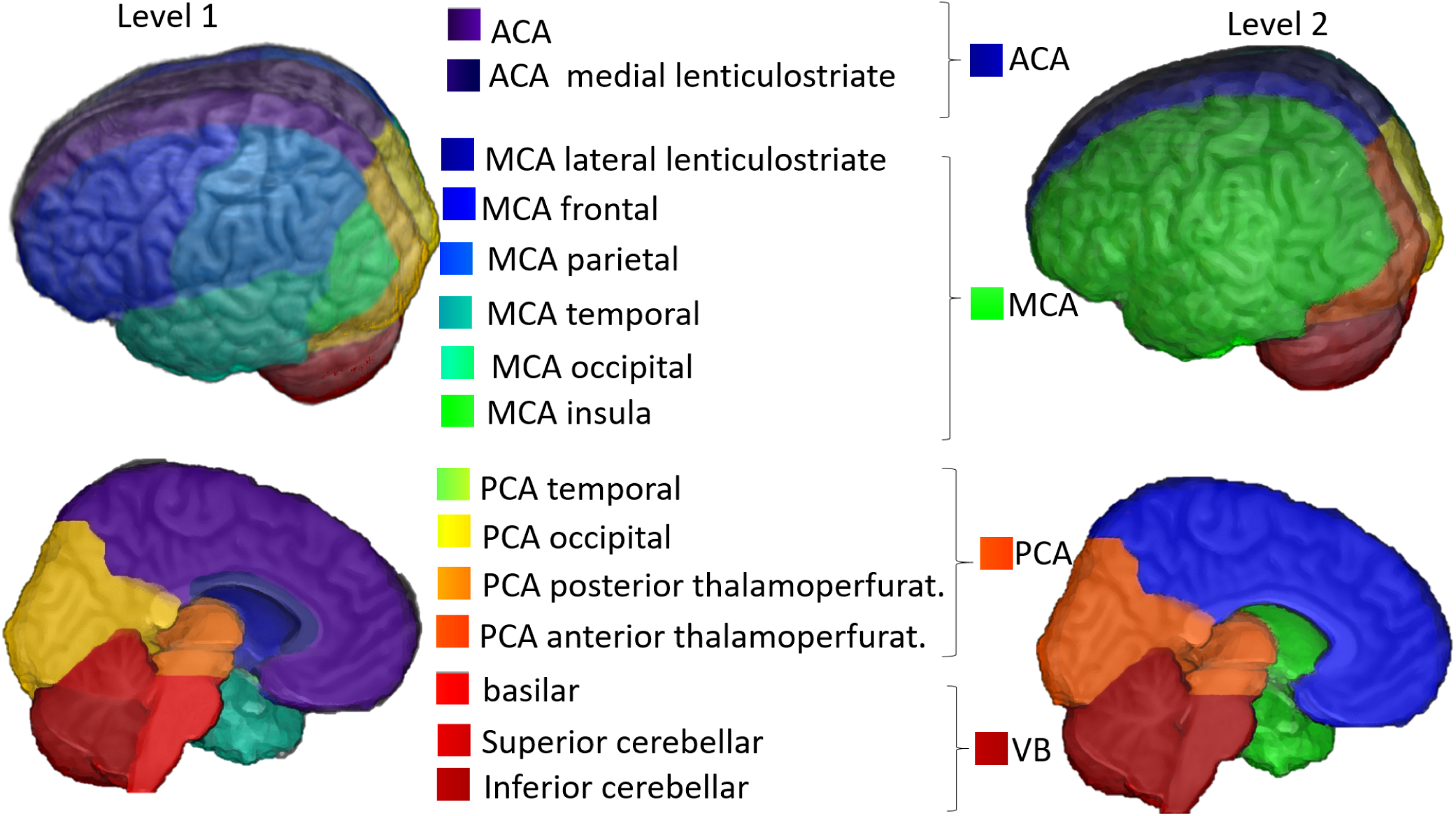
3D reconstruction of the arterial atlas. The right column (“level 2”) represents the major vascular territories; the left column (“level 1”) shows subdivisions of biological importance, based on probabilistic maps and / or anatomical landmarks.

### 2.6 Atlas of Arterial Territories

Figures 7 A-C show the probabilistic maps of strokes in “sublevels” of MCA (lateral lenticulostriates, 7 A), PCA (posterior thalamoperforating, 7 B), and VB (anterior and posterior cerebellar, and basilar; 7 C). The sample size employed for each probabilistic map is summarized in Figure 1. Figure 7 D shows the border zone between the MCA-PCA and the MCA-ACA territories, represented by the ratio between the probabilistic maps. Values higher than 1 indicate that the given “border zone” voxel is more often affected by MCA strokes than by PCA or ACA strokes.

**Figure 7.**
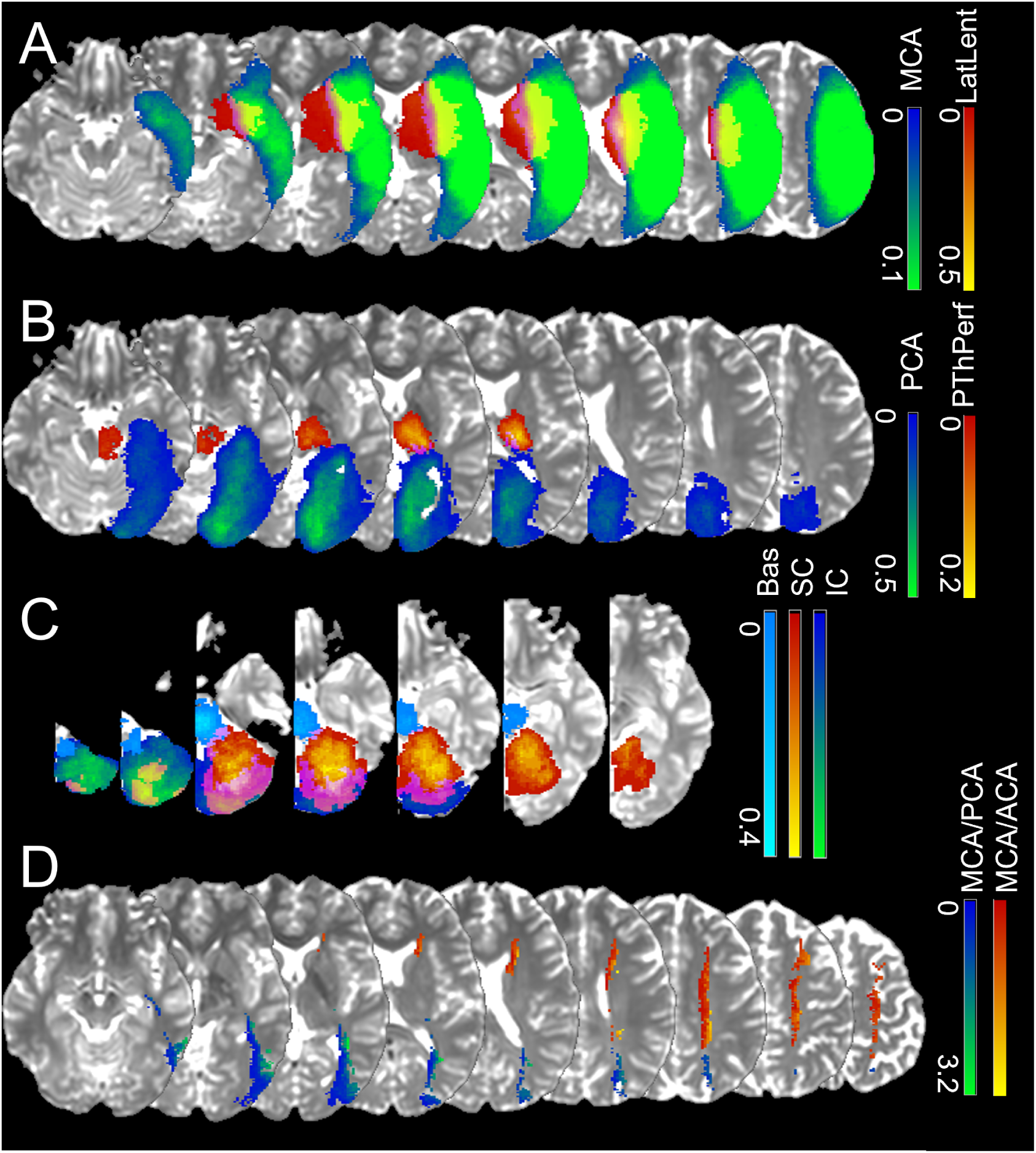
Probabilistic maps of strokes in “sublevels” of the major territories, with lesion affecting: (A) territories of lateral lenticulostriate arteries exclusively, or the “lobar” (cortex and adjacent white matter) MCA; (B) the territories of posterior thalamoperforating arteries exclusively, or the “lobar” (cortex and adjacent white matter) PCA; (C) the vertebro-basilar territory, composed of the anterior cerebellar, posterior cerebellar and basilar arteries. Color scales for each territory indicate the frequency of lesions. (D) shows the ratios between MCA-PCA (blue-green) and MCA-ACA (red-yellow) average probabilistic maps. Values higher than 1 indicate that the given “border zone” voxel is more often affected by MCA strokes than by PCA or ACA strokes.

Figure 8 is the 2D version of Figure 6 and shows our arterial territory atlas overlaid in T1-weighted axial slices of a brain in standardized MNI space. The bottom row shows the major arterial territories (ACA, MCA, PCA, VB); the top row represents the sublevels defined in the major territories by probabilistic maps (lateral lenticulostriate, posterior thalamoperforating, anterior and posterior cerebellar, and basilar strokes, as in Figures 7 A-C) or by prior knowledge of structural anatomy (medial lenticulostriate, anterior thalamoperforating, and “lobar” segments).

**Figure 8.**
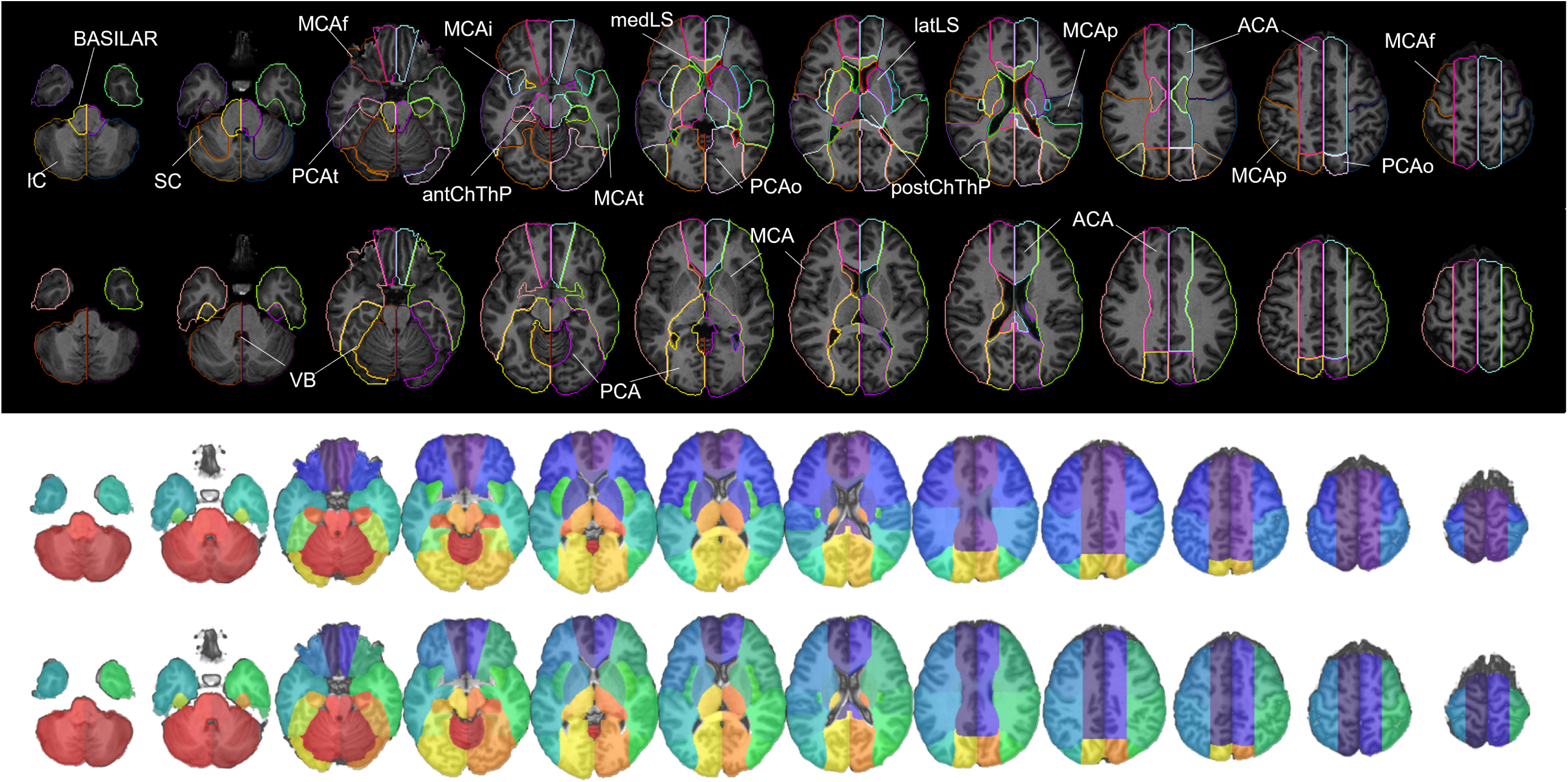
Digital arterial atlas overlaid on axial T1WIs of the template in standardized space (MNI). At the bottom of the superior panel, the major arterial territories: anterior (ACA), middle (MCA), and posterior (PCA) cerebral arteries, and vertebro-basilar (VB). At the top of the superior panel, subsegments of ACA (frontal (ACAf) and medial lenticulostriate (medLS)), MCA (frontal (MCAf), parietal (MCAp), temporal (MCAt), occipital (MCAo), insula (MCAi) and lateral lenticulostriate (latLS)), PCA (temporal (PCAt), occipital (PCAo), Posterior Choroidal and Thalamoperforating (postChThp), Anterior Choroidal and Thalamoperforating (antChThp)), and VB (inferior cerebellar (IC), superior cerebellar (SC), and basilar). For a clearer illustration, the bottom panel shows the same definitions (major territories at the bottom and subsegments at the top), overlaid in the template brain. The color code follows Figure 6, which corresponds to the atlas 3D reconstruction.

The sharp boundaries of our atlas qualitatively agree with the current knowledge of vascular territories, obtained by chemical injection in cadavers and *in vivo* image methods, as discussed in the Technical Validation. The quantitative level of agreement was also reasonable, although measured indirectly (see Technical Validation), since there is no counterpart deformable 3D atlas available for direct comparison.

### 2.7 Limitations

Although we used data from a certified “Comprehensive Stroke Center”, whose population reflects the profile of the national population with stroke, and scans with great technical heterogeneity (collected along 10 years in seven scanners from four manufacturers, and with dozens of different protocols), a regional bias may exist in the data. We expect the application of this Atlas to data from other centers and its potential enrichment with such data, as well as the users’ feedback through the public platform in which the Atlas is shared (described in “Data records”) will help us to identify and correct possible biases.

A limitation of this study is that the classification of vascular territories was based on expert description, rather than angiography (although 59% of the cases had MR angiography confirming the occlusion or stenosis of a single relevant large supratentorial artery and patent Carotids). The facts that we excluded “watershed”, cardioembolic, and “multifocal” strokes, and focused on reasonably large vascular areas help to ameliorate this issue. Nevertheless, we acknowledge the possible imprecision and the circular nature of the reasoning on using previous knowledge to create a tool to enrich such knowledge. While this seemed inevitable for the generation of this first version, we again hope that sharing data initiatives will enable us and others to accumulate data enough to create the next version of this Atlas based on an independent inclusion criterion.

Another limitation is that less stroke-prone territories (medial lenticulostriate and anterior thalamopefo-rating) were defined based in prior anatomical knowledge, rather than in probabilistic lesion maps (which would be unreliable given the present sample). We aim to redefine these boundaries using probabilistic maps, once we accumulate additional cases of strokes in these areas. Similarly, ACA strokes were less represented in our sample which might had affected the definition of border zones. While the BMM method was able to capture the correlation between voxels and variations in lesion volume and spatial features in each group, and helped to ameliorate the influence of the groups unbalance, the future inclusion of more ACA strokes will enable a better assessment of this territory.

Finally, we present probabilistic atlases calculated by two methods: averaging and BMM. There are numerous other strategies to generate atlases, each of them with specific advantages. It is unlikely that we or other groups can, alone, explore all the possible approaches. Therefore, sharing the individual images and the lesion maps, after fulfilling all the regulatory demands, is in our near future scope. This will enable researchers to create atlases and other tools that better fit the specific aims of their studies. For now, and despite all these limitations, the atlas of the arterial territories presented here may serve as a valuable resource for large-scale, reproducible processing and analysis of brain MRIs of patients with stroke, and other conditions.

## 3 DATA RECORDS

The atlas of the arterial territories is available in NITRC (https://www.nitrc.org/projects/arterialatlas). Any questions, concerns, or comments related to the atlas are welcome and can be publicly reported, so users are aware and we can resolve any issues in a timely manner. The images are in Montreal Neurological Institute (MNI) coordinates and the data format is “Nifti”, as recommended by the Brain Imaging Data Structure (BIDS). The description of the files is as follows:

1. ArterialAtlas.nii: Image defining 30 arterial territories and ventricles. The intensities of the parcels correspond to their labels IDs, listed in the lookup table “ArterialAtlasLabels.txt”, that accompanies the documentation. If visualized in 3D, the ArterialAtlas.nii corresponds to the left of Figure 6 of this paper; if visualized in 2D over anatomical slices, it corresponds to the top row of Figure 8 panels.
2. ArterialAtlas_level2.nii: The combination of ArterialAtlas.nii parcels in 4 major territories (ACA, MCA, PCA, VB). Again, the intensities of the parcels correspond to their labels, listed in the ArterialAtlasLabels.txt. If visualized in 3D, the ArterialAtlas_level2.nii corresponds to right column of Figure 6 of this paper; if visualized in 2D over anatomical slices, it corresponds to the bottom row of Figure 8 panels.
3. ArterialAtlasLabels.txt: is the “lookup table”. It contains the labels (descriptive and acronyms) for the regions defined in ArterialAtlas.nii and ArterialAtlas_level2.nii. The intensity of the region in the images corresponds to the label ID in the lookup table.
4. ProbArterialAtlas_average.nii: 4D image of the arterial territory maps, calculated by averaging of lesion masks, as detailed in the section 2.3. Each dimension represents the probability of a voxel to belong to a certain vascular territory (ACA, MCA, PCA, VB, respectively). If visualized in 2D over anatomical slices, it corresponds to the top row of Figure 3.
5. ProbArterialAtlas_BMM.nii: 4D image of the arterial territory maps, calculated by the BMM method, as detailed in the section 2.4. Each dimension represents the probability of a voxel to belong to a certain vascular territory (ACA, MCA, PCA, VB, respectively). If visualized in 2D over anatomical slices, it corresponds to the bottom row of Figure 3.
6. BorderZone_ProbAve.nii: 4D image showing the ratio of average probability maps, MCA / ACA and MCA / PCA. If visualized in 2D over anatomical slices, it corresponds to Figure 7D.
7. TerritoryVoxels_BMM.nii: 3D image showing the assignment of voxels to major vascular territories, calculated through the variation in BMM *µ*_*max*_, as described in section 2.4. The image intensities are: 1. ACA, 2. MCA, 3. PCA, 4. voxels attributed to more than one vascular territory, 5. voxels not attributed to any vascular territory. If visualized in 2D over anatomical slices, it corresponds to the top row of Figure 4.

## 4 TECHNICAL VALIDATION

### 4.1 Lesion Delineation

Although there is no perfect method for defining the stroke core, we chose to use DWI and Apparent Diffusion Coefficient maps (ADC), based on the fact that DWI is the most informative and most common sequence performed for acute stroke detection. Likewise, prior acute stroke studies and trials have defined lesions in DWI, so our data will be broadly comparable to those investigations. The delineation of the stroke core was defined in the DWI by 2 experienced evaluators and revised by a neuroradiologist until reaching a final decision by consensus. The evaluators looked for hyper intensities in DWI and / or hypo intensities (<30% average brain intensity) in ADC. Additional contrasts were used to rule out chronic lesions or microvascular white matter disease. A “seed growing” tool in ROIEditor (MRIStudio.org) was often used to achieve a broad segmentation, followed by manual adjustments. In a subset of 130 training cases (random selected 10% of cases) traced twice in a two-week interval, the index of agreement, DICE, was 0.76±0.14, inter-evaluator, and 0.8±0.13, intra-evaluator. Although this is a satisfactory level of agreement (Dice values range from 0 to 1; where 1 indicates total agreement), it demonstrates that human segmentation has suboptimal reproducibility, even when performed by experienced, trained evaluators, reinforcing the importance of the consensus agreement and the multiple revisions performed.

### 4.2 Image Mapping

The parameters for the non-linear deformation were chosen to maximize the accuracy on mapping structures such as the ventricles and the brain contour. For the linear registration we used the default setting of the affine transformation in BROCCOLI RegisterTwoVolumes function^28^. As for the non-linear transformation, there are two important parameters, sigma and iterationsnonlinear (N_iter), to determine the smoothness of the displacement field and the number of iteration steps. To find the optimal sigma and N_iter, we “hyperparameter-searched” all the combinations of sigma in [1,3,5] and N_iter in [10, 15, 20] to register a set of 470 subjects with “not visible” strokes to the template. We evaluated the deformation results for each combination of parameters across all subjects, as follows:

a. Three regions of 5 voxels bandwidth were defined in the template: the outside strip of the brain mask (OSBM), the inside strip of the brain mask (ISBM), and the outside strip of the lateral ventricles (OSLV), as shown in Figure 9, top.
b. The ratio of the mis-deformed voxels in OSBM, ISBM and OSLV for each subject, defined as *γ*OSBM, *γ*ISBM, and *γ*OSLV, was calculated as follows:

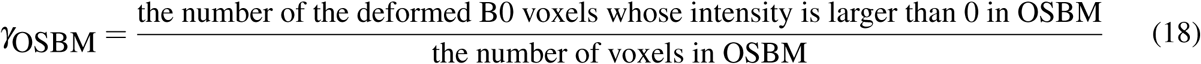

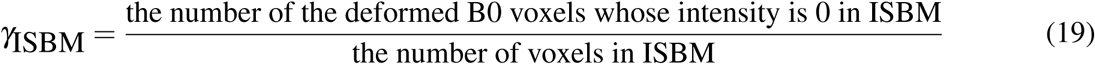

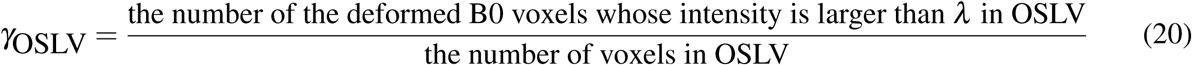

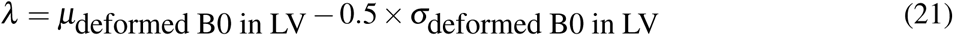 *γ*OSBM indicates the ratio of the deformed B0 voxels aligned outside the template brain mask and *γ*ISBM indicates the ratio of the background voxels aligned inside the template brain mask. High *γ*OSBM or *γ*ISBM mean the global deformation on B0 boundary is not good enough. *γ*OSLV indicates the ratio of voxels from a subject’s deformed ventricles that exist outside the template lateral ventricle boundaries. High *γ*OSLV is common in this population since aged subjects’ lateral ventricles are usually larger than the template’s lateral ventricles and indicated the need for local deformation.
c. The optimal parameters were sigma=3 and N_iter=15 (Figure 9, bottom). The registration with these parameters achieved better global deformation around the brain mask and comparable (to sigma=1 and N_iter=15) local deformation around the lateral ventricles.

**Figure 9.**
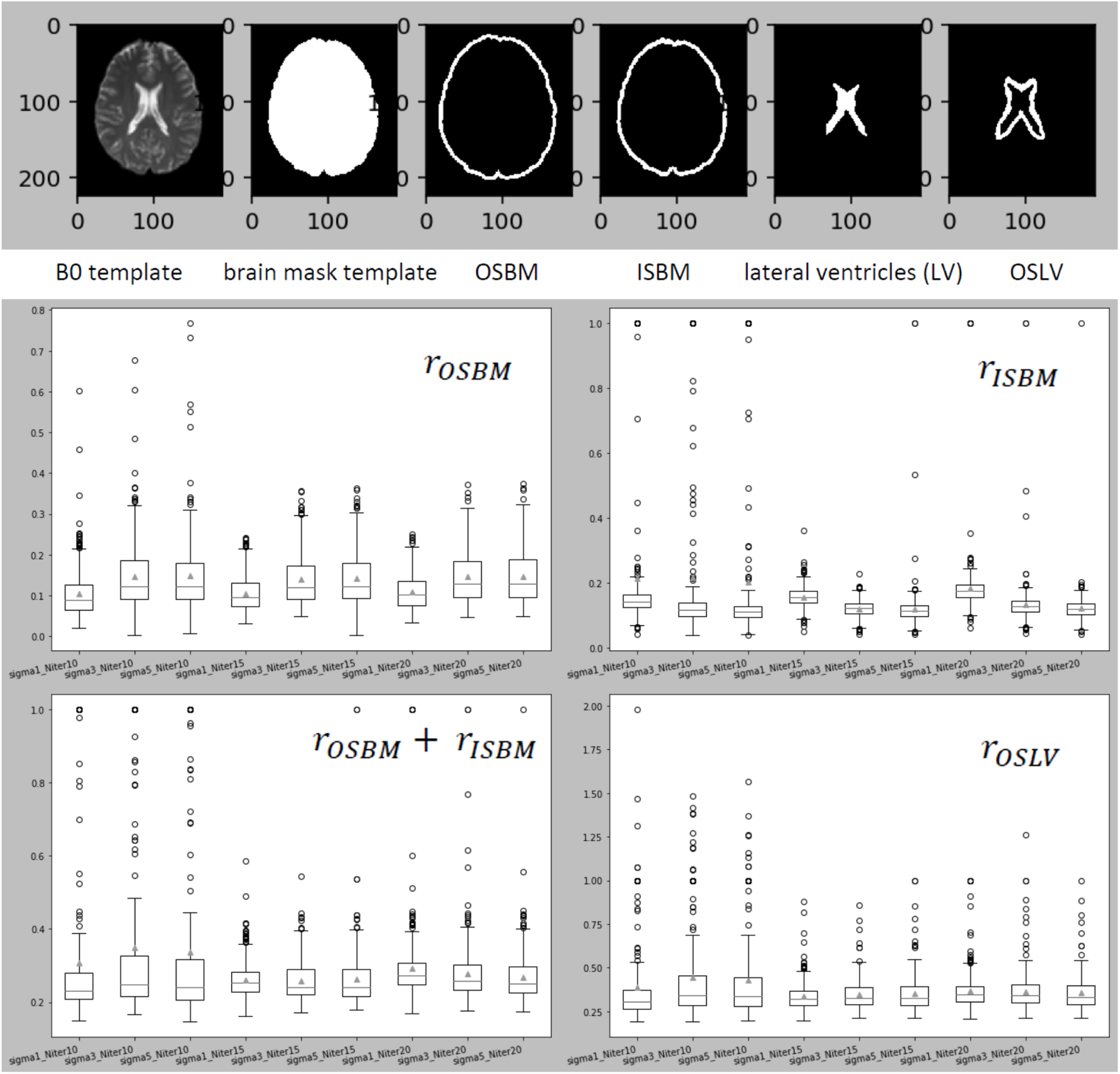
The top figure illustrates the three regions of 5 voxels bandwidth were defined in the template: the outside strip of the brain mask (OSBM), the inside strip of the brain mask (ISBM), and the outside strip of the lateral ventricles (OSLV). The bottom boxplots figure illustrates the statistics of *γ*OSBM, *γ*ISBM, and *γ*OSLV for all parameter’s combinations tested,

Visual quality control followed parameter optimization. The deformation matrix of each image was then applied to the respective stroke mask using near neighbor interpolation to preserve the binary nature. Images of patients with strokes in the right hemisphere were flipped along the x-axis so that all the stroke masks were considered in the left hemisphere in the standard space.

### 4.3 Clusters within arterial territories calculated by BMM

The probabilistic maps *µ*_*k*_ of ACA, MCA, PCA, and VB for the clusters of lesions in major vascular territories are shown in Figure 10. The size, prior probability, and mean volume of lesions in each cluster is shown in Table 2. Compared to the calculation by simple average, the range of probabilities calculated by BMM is higher, particularly in peripheral areas of each territory. The MCA clusters 1, 3, and 4 are anatomically similar to the frontal, parietal areas, and lateral lenticulostriate territory; the PCA clusters 1 and 2 are anatomically similar to the occipital area and the posterior Choroidal Thalamoperfurating territory; the VB clusters are anatomically similar to the inferior and superior cerebellar and basilar territories.

**Table 2.** The summary table of the *π*_*k*_, *N*_*k*_, and *E*[*V*_*k*_] for the BMM clusters

| Vascular Territories | ACA |  | MCA |  |  |  | PCA |  |  | VB |  |  |
| --- | --- | --- | --- | --- | --- | --- | --- | --- | --- | --- | --- | --- |
| Number of clusters | $K = 2$ | | $K = 4$ | | | | $K = 3$ | | | $K = 3$ | | |
| The $k^{th}$ Cluster | $\mu_1$ | $\mu_2$ | $\mu_1$ | $\mu_2$ | $\mu_3$ | $\mu_4$ | $\mu_1$ | $\mu_2$ | $\mu_3$ | $\mu_1$ | $\mu_2$ | $\mu_3$ |
| $\pi_k$ | 0.759 | 0.241 | 0.157 | 0.096 | 0.140 | 0.607 | 0.755 | 0.128 | 0.117 | 0.157 | 0.122 | 0.721 |
| $N_k$ | 22 | 7 | 120 | 73 | 107 | 463 | 271 | 46 | 42 | 23 | 18 | 106 |
| $E[V_k]$ in ml | 6.17 | 29.23 | 63.35 | 239.39 | 47.24 | 5.95 | 2.58 | 46.51 | 10.48 | 39.90 | 9.20 | 2.03 |

**Figure 10.**
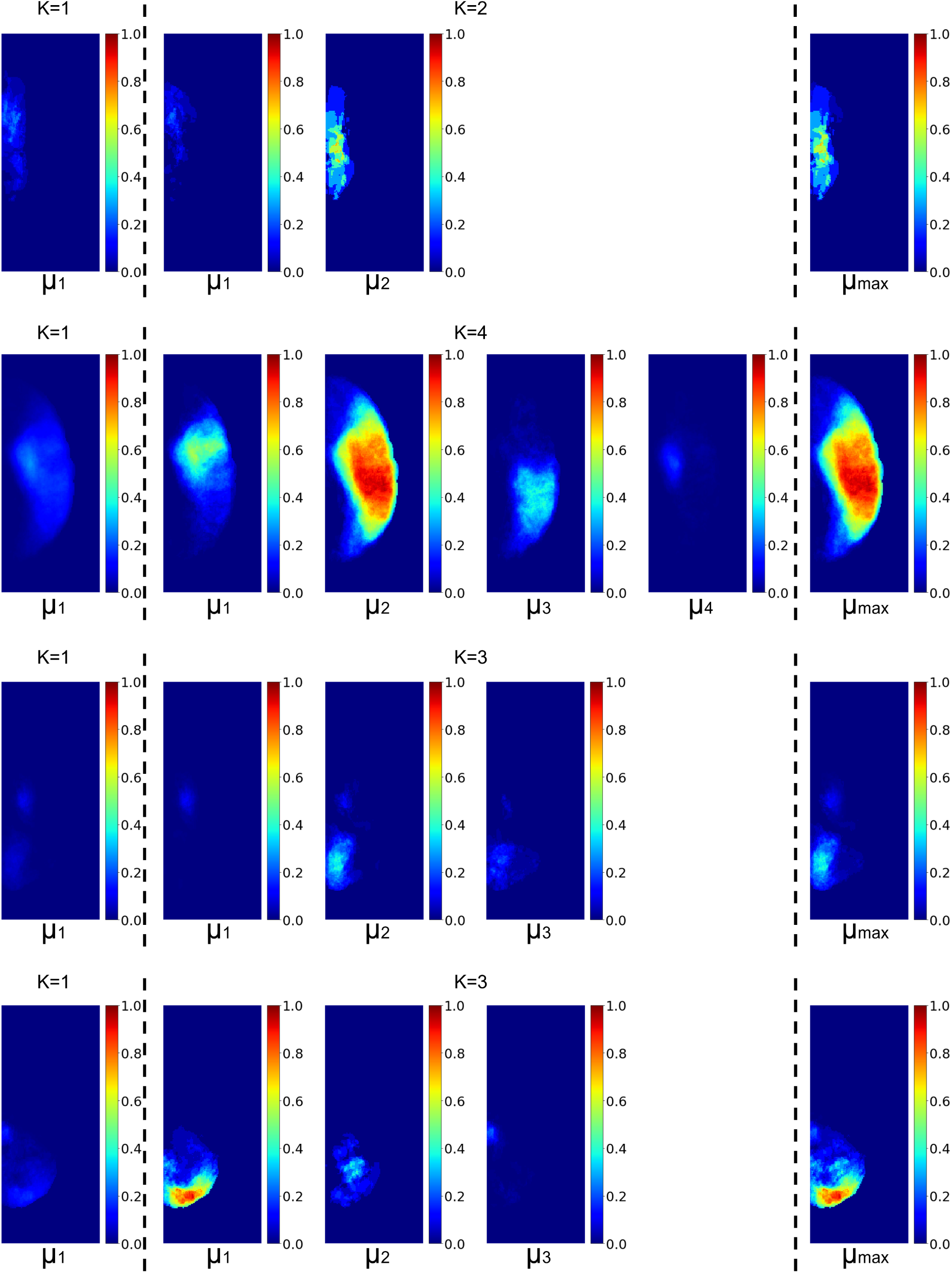
The probabilistic maps *µ*_*k*_ of ACA, MCA, PCA, and VB (rows, from top to bottom), for the average method, *K* = 1 (left panel), and for the clusters generated with BMM (middle panel, between the traced lines). The right panel shows the final *P*_*s*_(*X*_*j*_) = *µ*_*max*_ for each territory, obtained with the BMM model.

### 4.4 Qualitative Comparison to Previous Atlases

Figure11 shows the major arterial territories as defined in the present study overlaid in the left hemisphere, and an anatomical atlas (AAL) overlaid in the right hemisphere. The present atlas vastly agrees with Tatu et. al^6^ (sections I-XII, figure not reproduced here due to license protections). Two noticeable differences are: 1) The lateral extension of PCA. According to Tatu et. al, the PCA territory extends to most of medial and superior occipital gyri, while we found these areas are partially in the MCA territory, in agreement with recent stroke and perfusion-based MRI studies^17–19^. 2) The edges of the ACA-MCA-PCA territories, in the transition of the parietal and occipital lobes. While Tatu et. al^6^ attributed Pre Cuneus and Superior Parietal to ACA mostly, we found that Superior Parietal is mostly in MCA and PCA territory, reducing the lateral superior limits of ACA, and Pre Cuneus is partially in the PCA territory, expanding the superior limits of PCA. This is in alignement to the findings of van Laar et. al^14^ in cadavers and, more recently, Phan et. al^19^ in perfusion MRI.

Finally, in Figure12, we broadly illustrate the agreement of the present Atlas with digital reconstructions of brain arterial vasculature from MR angiography^29^, and with estimations of perfusion territories obtained by the coupling between arterial blood flow and tissue perfusion^6, 30^

**Figure 11.**
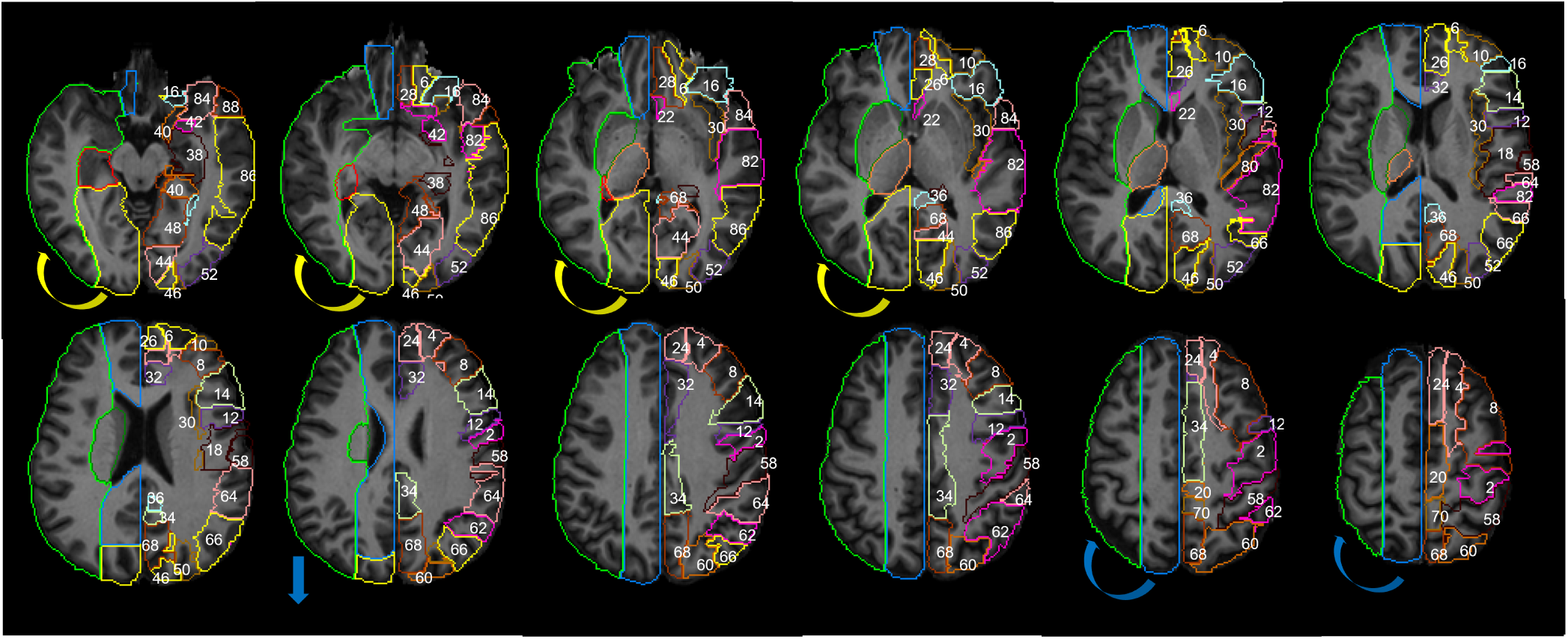
The JHU_MNI_SS template^25^ and the Automated Anatomical Atlas, AAL^31^, are used to display anatomic information in the supratentorial brain: z = 58, 64, 70, 76, 82, 88, 94, 100, 106, 112, 118, 124 mm. The numbers in the left hemisphere represent the following areas, from the AAL: 2. precentral, 4. dorsolateral superior frontal (GFs), 6. orbital superior frontal (GFso), 8. middle frontal (GFm), 10. orbital middle frontal (GFmo), 12. opercular inferior frontal (GFio), 14. triangular inferior frontal (GFit), 16. orbital inferior frontal (GFi), 18. rolandic operculum, 20. supplementary motor, 22. olfactory cortex, 24. medial superior frontal (GFds), 26. medial orbital superior frontal (GFdo), 28. rectus gyrus, 30. insula, 32. anterior cingulate and paracingulate, 34. median cingulate and paracingulate, 36. posterior cingulate, 38. hippocampus, 40. parahippocampus, 42. amygdala, 46. cuneus, 48. lingual (Gol), 50. superior occipital (GOs), 52. middle occipital (GOm), 54. inferior occipital (GOi), 56. fusiform (FG), 58. postcentral, 60. superior parietal (GPs), 62. inferior parietal (Gpi), 64. supramarginal (GSM), 66. angular (GA), 68. precunues (PreCu), 70. Paracentral lobule, 80. Hershel gyrus, 82. superior temporal gyrus (GTs), 84. superior temporal pole (GTps), 86. middle temporal gyrus (GTm), 88. middle temporal pole (GTpm), 90. inferior temporal (GTi). The colors in the right hemisphere are the major arterial territories as define by the present Atlas. The color code follows sections I-XII in Tatu et. al^6^, for easy comparison: ACA in blue, MCA in green, PCA in yellow. The arrows represent the main differences from Tatu et. al: the lateral inferior extension of PCA (yellow) and the lateral superior extension of ACA (blue) are larger in Tatu et. al Atlas, compared to ours.

**Figure 12.**
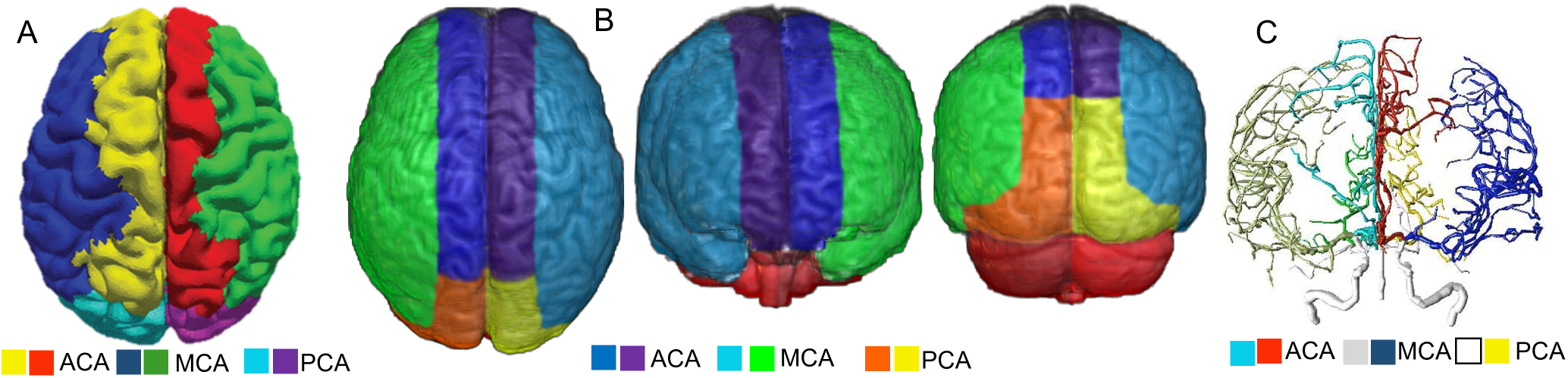
Qualitative comparison of our Atlas (B, axial superior, coronal anterior and posterior views) with estimations of perfusion territories obtained by the coupling between arterial blood flow and tissue perfusion^30^ (A, axial superior view), and with a digital reconstruction of brain arterial vasculature from MR angiography with BRAVA^29^ (C, coronal anterior view)

### 4.5 Quantitative Comparison to Previous Atlases

Since we present the first 3D deformable brain atlas of arterial territories, a direct quantitative comparison with counterparts is not possible. Therefore, we show an indirect comparison using the certainty index (CI), as^18^. The CI was calculated as

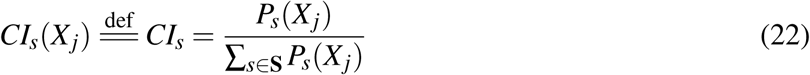

The CI can be interpreted as a conditional probabilistic map of voxel *X*_*j*_ belonging to *s* group given the likelihood of each voxel, *P*_*s*_(*X*_*j*_). The accuracy of CI depends on the sample size of each territory, the balance of sample sizes among groups, and the variation of lesions within each territory. As the average method, it ignores the correlation between voxels and the variations in volume and spatial features in each group.

Table 3 summarizes the CIs of our average probability maps. This table is organized as the supplemental table 2 in^18^ for easy comparison. Our CIs and those in Kim et. al are greatly aligned except in two areas. First, we obtained higher CIs in the occipital superior gyri in the PCA group. As our PCA sample size is significantly larger than that in Kim et. al, we have a higher level of confidence to attribute a greater portion of this area to PCA. This is also in agreement with digital maps of PCA infarcts^1, 19^ and the acknowledgement in Kim et. al that the PCA territory as they found is more constrained to medial areas than expected. Second, we obtained high CIs in the frontal superior and middle orbital areas in the MCA group. Still, we attribute great portions of these areas to ACA, in our sharp-boundaries Atlas. This is because, given the highly dependence of CIs on the sample size, on the balance among groups size, and our small ACA sample, we opted to relay on the projection of border zones and on the BMM “territory-claimed” voxel maps to trace the atlas sharp borders, rather than in CIs. This is in agreement with previous studies on different data modalities, including Tatu et. al^6^, that attributed those frontal areas to both ACA and MCA territories. As mentioned in the limitation section, the increase of the ACA sample will enable a better assessment of this territory.

**Table 3.**
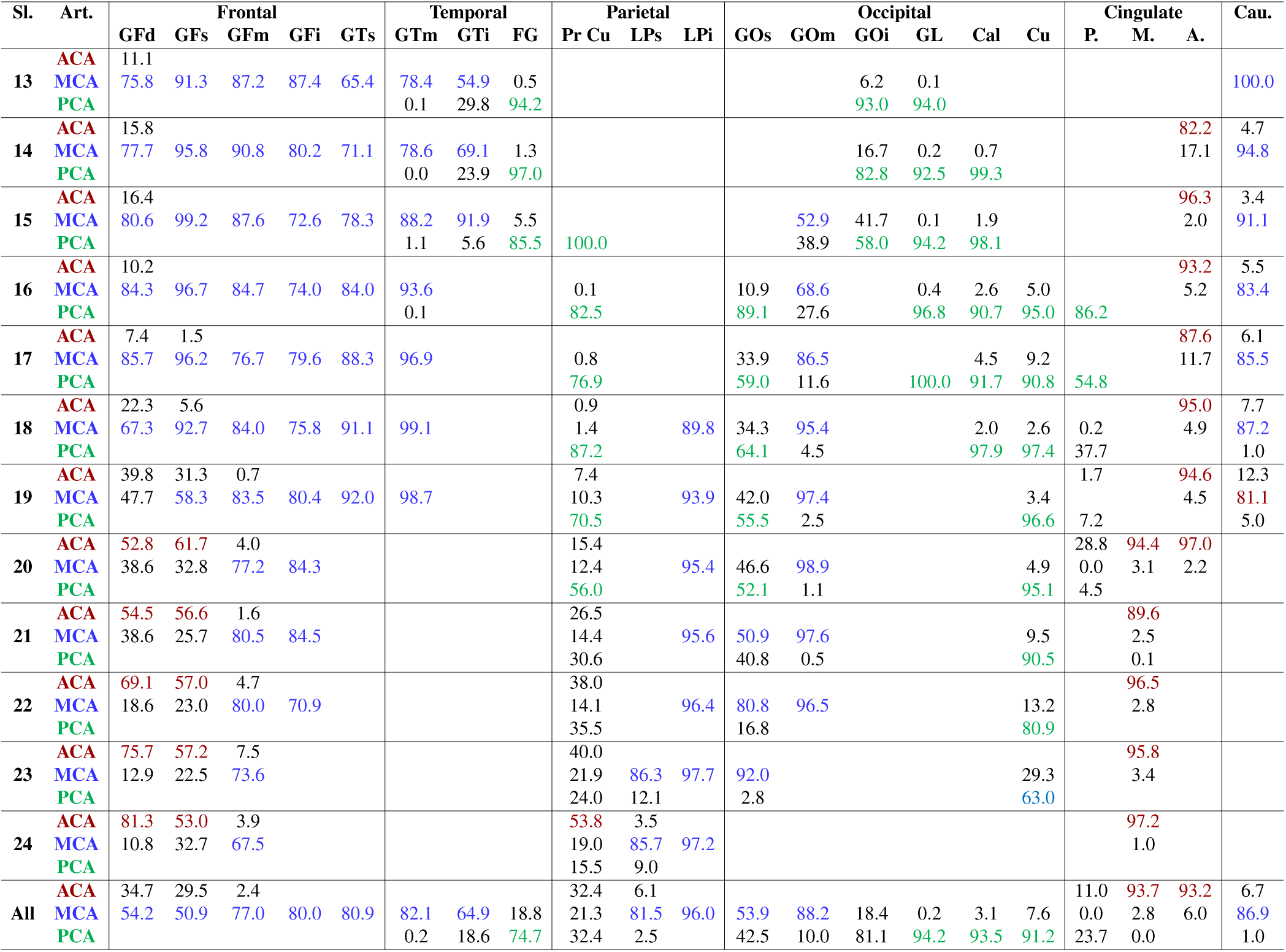
Mean Certainty Index, CI, in percentage, for diverse ROIs and their attribution to a Vascular Territory. This table is organized as in Kim et. al^18^, for easiness of comparison. The ROIs are defined by the Automated Anatomical Atlas, AAL^31^. CIs higher than 50% are colored as in^18^. Abbreviations: Sl. no, slice nuber; Art, artery; P, posterior; M, middle; A, anterior; Cau, caudate. For the abbreviations that follow, see caption of Figure 11. GFd includes GFds and Gfdo; GFs also includes Gfso; GFm also includes GFmo; GTs also includes GTps; GTm also includes GTpm; GFi includes GFio, GFit, and GFio; LPi also includes GSM and GA. The slice numbers (13-24) are compatible with those used by^18^ to calculate CIs, and with the sections I-XII in^6^. These slices correspond to the template (JHU_MNI_SS)^25^ coordinates 56-127 mm, grouped by 6, from bottom to top.

## Funding

This research was supported in part by the National Institute of Deaf and Communication Disorders, NIDCD, through R01 DC05375, R01 DC015466, P50 DC014664 (AH), and the National Institute of Biomedical Imaging and Bioengineering, NIBIB, through P41 EB031771 (MIM, AVF).

## Author contribution

AVF and CL conceived and designed the study, analyzed, and interpreted the data, drafted the work. JH, XX, GK analyzed the data. AEH, SS, EM acquired part of the data and substantially revised the draft. MIM revised the draft.

## Competing interests

Michael I. Miller owns “AnatomyWorks”. This arrangement is being managed by the Johns Hopkins University in accordance with its conflict-of-interest policies.

## Notes

https://www.nitrc.org/projects/arterialatlas/

### Summary of Updates

This version of the manuscript has been revised to include an extra calculation of probabilistic maps based on Bernoulli Mixed Models

